# Optimization of a Human Anti-polio Monoclonal Antibody as a Potential Therapeutic Modality

**DOI:** 10.64898/2026.09.20.753044

**Authors:** Shwu-Maan Lee, Christine Siska, Igor D’Angelo, Diana Kouiavskaia, Kutub Mahmood

## Abstract

**Background:** Vaccines have been an essential tool in bringing the world close to polio eradication, with over 99.9% of the global population free of poliovirus. No antiviral drugs or monoclonal antibody products, however, are currently licensed for treatment of polio. We previously isolated a human monoclonal antibody (9H2) with potent neutralizing activity against all three poliovirus serotypes. To advance 9H2 as a therapeutic candidate, we optimized its sequence to extend serum half-life and improve manufacturability.

**Methods:** A Multi-Attribute Method under stress conditions identified post-translational modification sites, and Abacus™ ranked sequence liabilities. Six amino acid substitutions were introduced into the variable regions, generating 23 combinatorial variants. Codon-optimized genes were synthesized and engineered into a human immunoglobulin G1 backbone containing crystallizable fragment (Fc) mutations (M428L/N434S) to extend serum half-life. Constructs were transfected into CHO-K1 cells to generate stable pools in 24-well plates. Protein A–purified antibodies were characterized using biophysical assays and an in vitro poliovirus neutralization assay.

**Results:** All variants retained high in vitro neutralizing activity against poliovirus. A lead candidate was selected based on integrated assessment of biophysical properties across eight assays and expression yield.

**Conclusions:** Structure-guided engineering and experimental evaluation enabled optimization of the 9H2 antibody sequence and identification of a lead candidate suitable for clinical manufacturing as a potential anti-poliovirus immunotherapeutic agent.

## BACKGROUND

Poliomyelitis remains one of the most intensively studied viral diseases and the subject of a sustained global effort towards eradication of polio. The causative agent, poliovirus (a positive-sense single strand RNA Picornaviridae enterovirus), is transmitted primarily via the fecal-oral route and, in some instances, can cause paralytic disease by infecting motor neurons.

The Global Polio Eradication Initiative (GPEI) has reduced cases of wild poliovirus (WPV) by more than 99.9 % since its launch in 1988, with success due, in large part, to the widespread use of live attenuated oral polio vaccine (OPV) and inactivated poliovirus vaccine (IPV). Two of the three historically recognized wild serotypes, wild poliovirus type 2 (WPV2) and WPV3, have been certified eradicated. WPV1 cases are restricted to Afghanistan and Pakistan. In addition, outbreaks due to circulating vaccine-derived polioviruses (cVDPVs) are predominantly type 2 virus, and types 1 and 3 viruses continue to be detected in different regions of the world with suboptimal immunity driven by genetic reversion of the attenuated OPV strains in under-immunized populations [1,2].

Extensive analyses of eradication efforts highlight the difficulty of fully interrupting poliovirus transmission. Key challenges include gaps in routine immunization coverage, limitations in surveillance systems, and the ongoing epidemiologic risk posed by stool shedding of cVDPVs and VDPV in immunocompromised individuals (iVDPVs), which may persist for several years in individuals with primary immunodeficiency disorders [1]. Vaccination remains the central strategy for polio control and eradication. New generation oral vaccine, the genetically stabilized novel OPV2 (nOPV2), has been deployed for cVDPV2 outbreak response use since 2021 [3].

Despite these successes, no therapeutic antiviral agent or antibody has been approved that directly targets poliovirus replication or pathogenesis after infection. Antiviral drug candidates (e.g., small molecule capsid binders) are being investigated in preclinical and early clinical studies; however, toxicity and resistance are concerns [4].

To address this deficiency, interest has grown in the development of monoclonal antibodies (mAbs) with potent neutralizing activity against polioviruses. A therapeutic human mAb could play a role in halting excretion in asymptomatic carriers and could be used, in combination with vaccines and/or antiviral drugs, to protect poliovirus-exposed individuals. Cross-neutralizing mAbs would be particularly useful because they would reduce the number of mAbs needed to create a comprehensive polio outbreak control therapeutic strategy for types 1, 2 and 3 polioviruses [5].

The human mAb 9H2, generated from a vaccinated donor, exhibits potent and uniquely broad neutralizing activity against all three poliovirus serotypes. It also demonstrates broad cross-neutralizing activity against a diverse panel of wild and vaccine derived polioviruses in clinical isolates. These factors support its potential as a broadly active therapeutic candidate [6].

Recent cryo-electron microscopy (cryo EM) structural studies show that mAb 9H2 binds at the 5-fold axis of the polio capsid within the conserved canyon region that overlaps with the poliovirus receptor (CD155) binding site. This binding prevents the receptor from virus attaching by both competing with the soluble receptor (PVR) and physically blocking access to binding site. This antibody binding mode is the same across serotypes 1–3, which explains the cross-reactive serotype neutralization without inducing major capsid changes, and makes 9H2 a strong candidate for development as a broadly active antiviral biologic for therapeutic use with low likelihood of simple escape mutants [7]. Further structure studies show that 9H2 shares capsid contacts with the receptor and blocks receptor binding, leading to virus particle neutralization and poliovirus entry inhibition into host cells [8].

To advance mAb 9H2 as a polio therapeutic modality, we optimized the antibody sequence to improve its serum half-life and manufacturability. These variants were then tested in the poliovirus neutralization assay to confirm their biological function.

## METHODS

### Monoclonal Antibodies Used in This Study

Human monoclonal antibodies (mAbs) were generated from peripheral blood of volunteers who received a booster dose of IPV eight days prior to sampling. CD27⁺ memory B cells were fused with the B5-6T myeloma cell line to produce hybridomas. Supernatants were screened by enzyme-linked immunosorbent assay (ELISA) for immunoglobulin G (IgG) binding to Sabin poliovirus types 1, 2, and 3, followed by microneutralization assays [6].

mAb 9H2, which exhibited broad neutralizing activity against all three poliovirus serotypes (Sabin and wild-type strains), was selected for this study; this breadth suggests potential activity against cVDPVs, which retain conserved neutralizing epitopes. In addition to the 9H2 there were other serotype-specific mAbs—2D6 (type 1), 1B8 (type 2), and 6B5 (type 3)— generated and used as controls in neutralization assays to confirm serotype specificity [9].

### Multi-Attribute Mass Spectroscopy (MAM) to probe Post Translational Modification (PTM)

MAM analysis was conducted on parental mAb 9H2 (untreated control), high-pH-treated (pH 8.5, 5 days at 40 ℃) and oxidation-stressed (0.05% H_2_O_2_, 1 day at 40 ℃) samples. These samples were denatured, reduced, alkylated, buffer exchanged, and trypsin digested using the same methods as those described by Ogata et al [10], except that 5 mM Tris(2-carboxyethyl) phosphine was used as a reducing agent.

Peptides from tryptic digest were analyzed by LC-MS/MS on a Vanquish LC in line with an Exploris-240 mass spectrometer (Thermo). A Zorbax C18 300-SB column (Agilent) was used with formic acid and Acetonitrile gradient to separate the peptides [10]. Peptide sequences and PTMs were identified and quantified using Byos^TM^ software (Protein Metrics).

### Abacus^TM^ in Silico Analysis of Parental mAb GH2 to Identify Sequence Liabilities

mAb 9H2 was subjected to structural modeling using Abacus**^TM^**, an in silico tool developed in-house at Just-Evotec. The platform analyzes and ranks antibody sequences, identifies potential liabilities affecting developability and stability, and predicts remediation strategies using a machine learning algorithm trained on the Observed Antibody Space (OAS). The parental sequence underwent liability analysis using Abacus**^TM^** and hotspot positions were mapped onto the complex crystal structure (Protein Data Bank, ID 8E8Y [11]) to better understand their implications and possible stability issues. The structure was also used as input in the MOE 2024.02 software suite (Chemical Computing Group, Montreal, Canada) to calculate surface properties (i.e., surface patches of different nature). A set of 23 designs was generated combining optimized mutations on the VL and VH.

### Generating Variants in the Deep Well Plates for Analysis

Codon-optimized DNA sequences encoding the designed variable regions of the heavy and light chains were chemically synthesized. A 24th variant containing no amino acid changes in the variable regions, was generated as a comparator. These sequences were engineered into expression vectors containing a human IgG1 backbone with Fc mutations M428L/N434S (LS) to extend antibody serum half-life [12].

Plasmids were transfected by electroporation into Glutamine Synthetase (GS)-null CHO-K1 cells in quadruplicate. Following transfection, cells were seeded into selection medium (CD OptiCHO™, Thermo Fisher Scientific). Cell cultures were maintained under standard conditions and monitored for viability and cell density, with periodic medium exchange during recovery and outgrowth of stable pools.

From the 24 deep well plates containing the stable pools, cells were seeded into production medium for a 10-day fed-batch production assay. Antibody variants were then harvested, followed by purification via Protein A column chromatography for biophysical analysis and functional assays. The cell pools were frozen for future scale up.

### Biophysical Analysis of the Variants

Eight biophysical methods were used to compare these variants. Table 1 outlines the name of these methods, the attribute measured, the characterization, the desired outcome, and the values indicating instability.

**Table 1:** The eight biophysical methods used to evaluate the variants.

| Method name | Attribute measured | Characterization | Desired outcome | Values that indicate instability |
| --- | --- | --- | --- | --- |
| Differential scanning fluorimetry (DSF) | Melting temperature | Lower melting temperature (T<sub>m</sub>) is indicative of decreased conformational stability | Increase in T<sub>m</sub> or formation of additional T<sub>m</sub>'s indicating increased domain stability | Weighted Shoulder Score (WSS) Below 20 |
| Thermal hold | Temperature induced aggregation | Define conditions for precipitation relating to potential destabilization during room temperature incubation | Increase in thermal stability indicated by the absence of precipitation | Absorbance at 350 nm is above 0.5 |
| Low pH aggregation | Low pH stability by measuring high molecular weight species | Molecules with an increase in high molecule weight following neutralization after low pH exposure may show increased aggregation during low pH viral inactivation | No significant increase in high molecular weight species following low pH exposure and neutralization | More than 10% aggregation after low pH exposure |
| Chemical unfolding with guanidium hydrochloride | Conformational stability | Molecules with an increased inflection point may show lower rates of aggregation during storage and are more conformationally stable | Increased inflection point compared to the parent molecule | Less than 2.1 M inflection point |
| Self-interaction nanoparticle spectroscopy (SINS)[15] | Tendency to self-associate via nanoparticle clustering | Define relative protein self-association to help identify potential for higher viscosity during concentration and problems with filterability | Lower interaction (minimal wavelength shift) | Wavelength Maximum Above 550 nm |
| Relative Solubility Analysis (RSA) | Polyethylene glycol based solubility analysis | Molecules with higher relative solubility may show lower rates of aggregation during storage | Increased solubility compared to parent molecule | 50% loss of protein occurring less than 7% PEG |
| Standup monolayer affinity chromatography (SMAC) on a Zenix column [16] | Colloidal stability | Molecules with reduced retention times may show increase solubility and lower rates of aggregation during storage | Decreased main peak retention time compared to parent molecule | Longer retention times; no established values correlate to instability at this time |
| Polyreactivity | Non-specific binding | Molecules that have the potential to non-specifically bind to an array of different antigens can potentially have a higher clearance rate. | Decreased non-specific binding compared to the parent molecule | Absorbance at 405 nm is above 1.5-2.0 |

### Poliovirus Neutralization Assay of the variants

The microneutralization test was performed as described in the World Health Organization (WHO) laboratory manual [13], with minor modifications. Twofold serial dilutions of the mAb variants and parental mAb 9H2 (starting at 10 µg/mL) were prepared in the maintenance medium (Dulbecco’s modified Eagle’s medium [DMEM] supplemented with 2% fetal bovine serum and 1% of antibiotic/antimycotic, Invitrogen) and mixed with equal volume of the challenge virus diluted to obtain 100 TCID_50_ (Tissue Culture Infectious Dose 50%) per well. The mixes were incubated at 36 °C for 3 hours. Hep-2C cells (ATCC) were added to the wells at the end of the incubation. Plates were incubated for 8–10 days at 36 °C in 5% CO₂. For assays using the S19 hyper attenuated strains, the incubation temperature was reduced to 33 °C [14]. Wells without cytopathic effect were counted, and poliovirus neutralization titers (defined as the dilution of mAb required to protect 50% of cell cultures) were calculated using the Kärber formula [13]. Neutralization titers were normalized to the original mAb concentration of 1 mg/mL.

The challenge viruses used in the neutralization assays were Type 1 strains Sabin NA4 and Wild Type 1 strain Mahoney; Type 2 strains S19-S2 [14] and S19-MEF1; and Type 3 strains Sabin NC2, S19-S3, and S19-Saukett. S19 strains were obtained from the National Institute for Biological Standards and Control (NIBSC, Potters Bar, UK). Serotype-specific mAbs (Type 1: 2D6, Type 2: 1B8, and Type 3: 6B5) [9] were included to confirm serotype specificity.

## RESULTS

### MAM Analysis

Post translational modifications of 9H2 were evaluated using MAM. Table 2 summarizes the results for untreated parental 9H2, as well as high-pH–treated and oxidation-stressed samples. Oxidative stress resulted in increased methionine and tryptophan oxidation. Three positions of interest in the heavy chain variable regions (HV) were identified as PTM hot spots, including one tryptophan in the framework-HV: W54 and two methionine in the complementarity-determining region (CDR): HV: M57 (CDR2) and HV: M136 (CDR3). Untreated sample also showed methionine oxidation greater than 10% in the HV:M57 position and it increased to greater than 95% under oxidation stress. The HV:M136 site showed no detected oxidation in the untreated sample, but an increase to 34.3% under oxidative stress conditions. Small increases (less than 10% total oxidation) were observed for tryptophan sites, including HV: W54 and HV: W115.

**Table 2:**
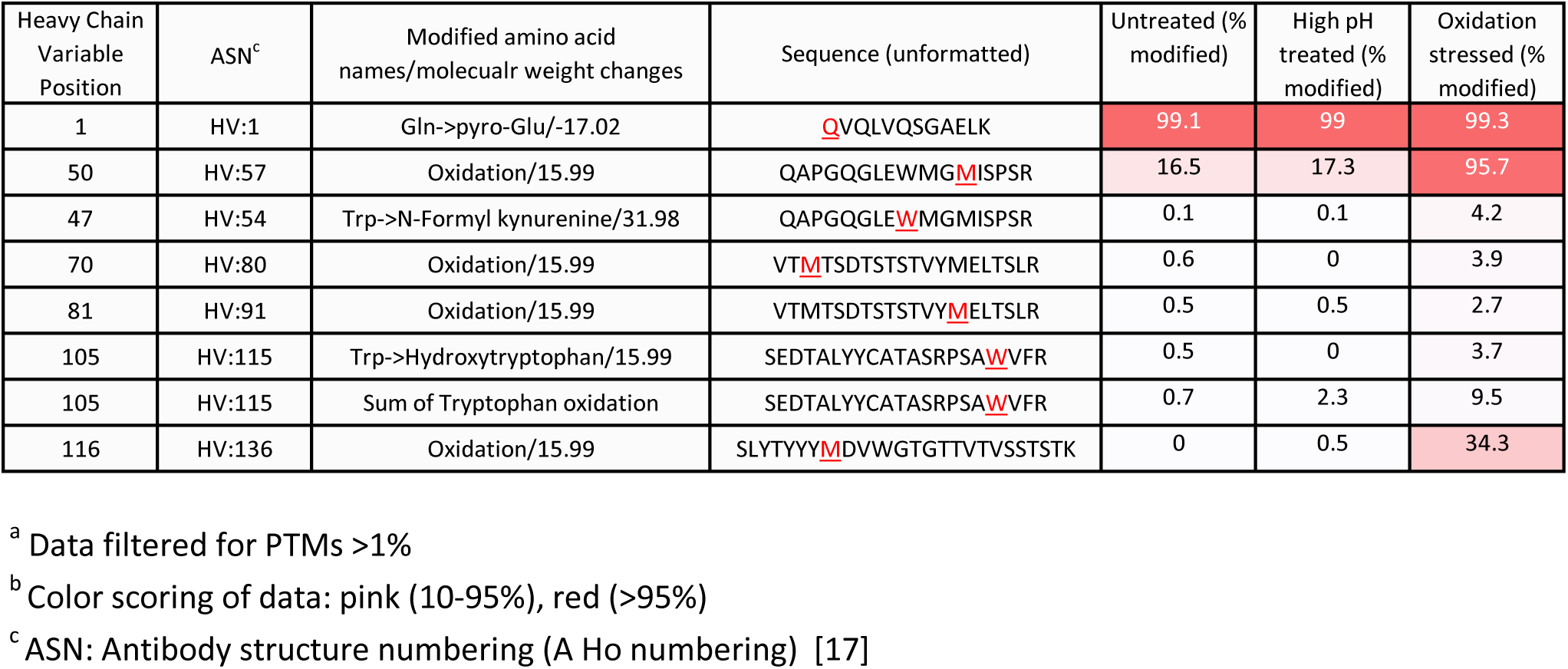
Post Translational Modifications (PTMs) from Multi Attributes Mass Spectrometry analysis^a,b^.

Further analysis of molecular structure indicated that these three hotspot residues were involved in the heavy chain and light chain variable regions interactions based on crystal structure [11], and they were not directly contacting the antigen. HV: M57 (CDR2) and HV: M136 (CDR3) were mostly buried and oxidation at these positions was considered a low developability risk. No mutation was pursued for HV: W54 based on MAM results with low percentage oxidation. No PTM of greater than 1% were observed in the light chain variable region.

### Abacus^TM^ Analysis of Parental mAb GH2

In silico analysis of the sequence identified several additional PTM hotspots, but MAM confirmed these levels as acceptable. Potential liabilities included isomerization at HV: D66 (CDR2) and Tryptophan oxidation at HV: W115 (CDR3); however, their abundance (less than 1% for D66 isomerization and W115 oxidation in the untreated sample) was not at a concerning level.

The analysis also indicated that a disrupted HV CDR3 salt bridge could be a potential issue. This might increase CDR3 flexibility, increase exposure of HV: W115 (oxidation risk), and worsen aggregation and off-target binding. In addition, the antibody contained two relatively large exposed hydrophobic patches (approximately 210 Å² and 150-200 Å²). Although such patches could increase aggregation risk, structural modeling based on PDB 8E8Z [11] suggested that these residues participated in antigen binding. Therefore, they were not altered during sequence optimization.

In summary, mAb 9H2 exhibited a localized, CDR-driven hydrophobic surface patch that may increase aggregation risk. Otherwise, it showed clean cysteine pairing, balanced charge, low immunogenicity, and acceptable physicochemical properties.

### Design of Variants

mAb 9H2 contained uncommon framework residue combinations that were not evolutionarily supported in human antibodies and might increase structural or developability risk according to RANDŸ (In-silico mutational tolerance analysis) statistical tolerance models. For variant design, most changes were conservative consensus corrections, but HV:108 represented a structurally meaningful salt-bridge repair opportunity likely to improve stability and reduce developability risk.

Based on Abacus**^TM^** predictive and structural analysis combined with results from MAM experiments, 23 variants were designed for the optimization of the mAb 9H2 (Table 3). These amino acid modifications were classified into the following three types and the variants consisted of at least one mutation:

1. Low-risk framework mutation: LV: G102A. This single mutation in light chain was applied to all variants
2. Medium risk structural refinements: HV: F21V, HV: Q24K, HV: F29Y, and HV: T141K. All these changes were in the framework of heavy chain.
3. High-impact stabilizer: HV: T108 R. It was expected to restore salt bridge.

**Table 3:** The comparator and the sequence changes for the 23 variants.

| Molecule | Antibody variant numbers | Light chain change | Heavy chain change |
| --- | --- | --- | --- |
| M-24210<sup>a</sup> | 9H2 | None | None |
| M-24211 | 9H2.001 | LV: G102A | HV:F21V |
| M-24212 | 9H2.002 | LV: G102A | HV:F29Y |
| M-24213 | 9H2.003 | LV: G102A | HV:T108R |
| M-24214 | 9H2.004 | LV: G102A | HV:F21V HV:Q24K |
| M-24215 | 9H2.005 | LV: G102A | HV:Q24K HV:F29Y |
| M-24216 | 9H2.006 | LV: G102A | HV:T108R HV:T141K |
| M-24217 | 9H2.007 | LV: G102A | HV:F21V HV:Q24K HV:T108R |
| M-24218 | 9H2.008 | LV: G102A | HV:Q24K HV:F29Y HV:T141K |
| M-24219 | 9H2.009 | LV: G102A | HV:F29Y HV:T108R HV:T141K |
| M-24220 | 9H2.010 | LV: G102A | HV:F21V HV:F29Y HV:T108R HV:T141K |
| M-24221 | 9H2.011 | LV: G102A | HV:F21V HV:Q24K HV:F29Y HV:T108R HV:T141K |
| M-24222 | 9H2.012 | LV: G102A | HV:F21V HV:Q24K HV:F29Y HV:T108R |
| M-24223 | 9H2.013 | LV: G102A | HV:F21V HV:Q24K HV:F29Y HV:T141K |
| M-24224 | 9H2.014 | LV: G102A | HV:F21V HV:Q24K HV:T108R HV:T141K |
| M-24225 | 9H2.015 | LV: G102A | HV:F21V HV:Q24K HV:T141K |
| M-24226 | 9H2.016 | LV: G102A | HV:F21V HV:F29Y HV:T108R |
| M-24227 | 9H2.017 | LV: G102A | HV:F21V HV:F29Y HV:T141K |
| M-24228 | 9H2.018 | LV: G102A | HV:F21V HV:T108R HV:T141K |
| M-24229 | 9H2.019 | LV: G102A | HV:Q24K HV:F29Y HV:T108R |
| M-24230 | 9H2.020 | LV: G102A | HV:Q24K HV:T108R |
| M-24231 | 9H2.021 | LV: G102A | HV:Q24K HV:T141K |
| M-24232 | 9H2.022 | LV: G102A | HV:F29Y HV:T108R |
| M-24233 | 9H2.023 | LV: G102A | HV:F29Y HV:T141K |

<sup>a</sup> M-24210 (the comparator) has no changes in the variable region, only with fragment crystallizable mutation to extend serum half-life

The 24^th^ variant containing no amino acid changes in the variable regions, only Fc mutation as the rest of the variants, the comparator, was denoted as M-24210.

### Generation of Variants and Evaluation of Their Growth Performance

A 10-day fed-batch production assay was carried out in 24-well plates. Cultures were harvested on day 10, antibody titers in the harvest were measured, and specific productivity (qP; picograms of antibody per cell per day) was calculated. All values are reported as averages of quadruplicate experiments (Figures 1). Several variants exhibited increased titers relative to the comparator (the 24th variant, M-24210), with most of the titers greater than 2.0 g/L. In contrast, the F29Y and T108R variants (M-24212 and M-24213; rightmost in Figure 1) showed markedly reduced titers, which correlated with lower cell growth and decreased viability during production.

**Figure 1.**
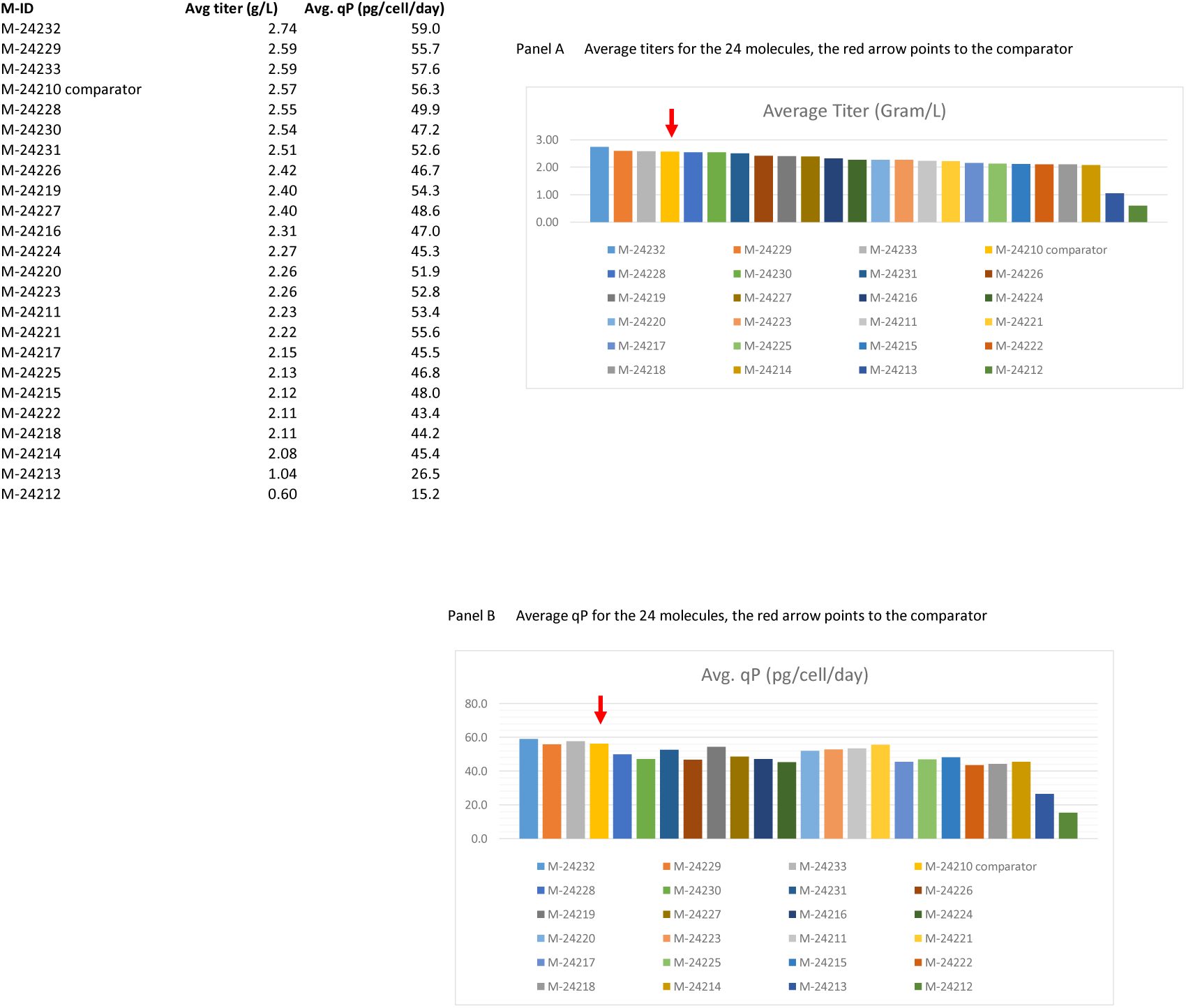
The average titers and qP (picograms of antibody per cell per day) for the variants expressed in the 24 well-plates (replicates of 4).

### Biophysical Analysis of the Variants

Eight biophysical methods were used to evaluate these variants. The compilation of these results is shown in Table 4, which includes the evaluated data color coded in green for acceptable, yellow for concerns, and red for unacceptable. Panel A presented data from all molecules. The comparator, M-24210, showed good biophysical characteristics and several variants slightly improved conformational stability (lower precipitation in thermal hold and less aggregation by size exclusion chromatography). The T108R mutation (included in 13 variants: M-24213, M-24216, M-24217, M24219-22, M-24224, M-24226, M24228-30, and M-24232) reduced colloidal stability based on Self-Interaction Nanoparticle Spectroscopy (SINS), which are unacceptable variants (color coded in red). In Panel B, the thirteen T108R variants were removed, and process performance data (titer and qP) were added. M-24227 showed the most improvement in thermal stability while maintaining high titer. M-24214 and M-24218 also showed improvement in thermal stability but had a slight reduction in titer as compared with the comparator M-24210. In Panel B, these three molecules are high-lighted in green. M-24227 is the top variant for future clinical development.

**Table 4:**
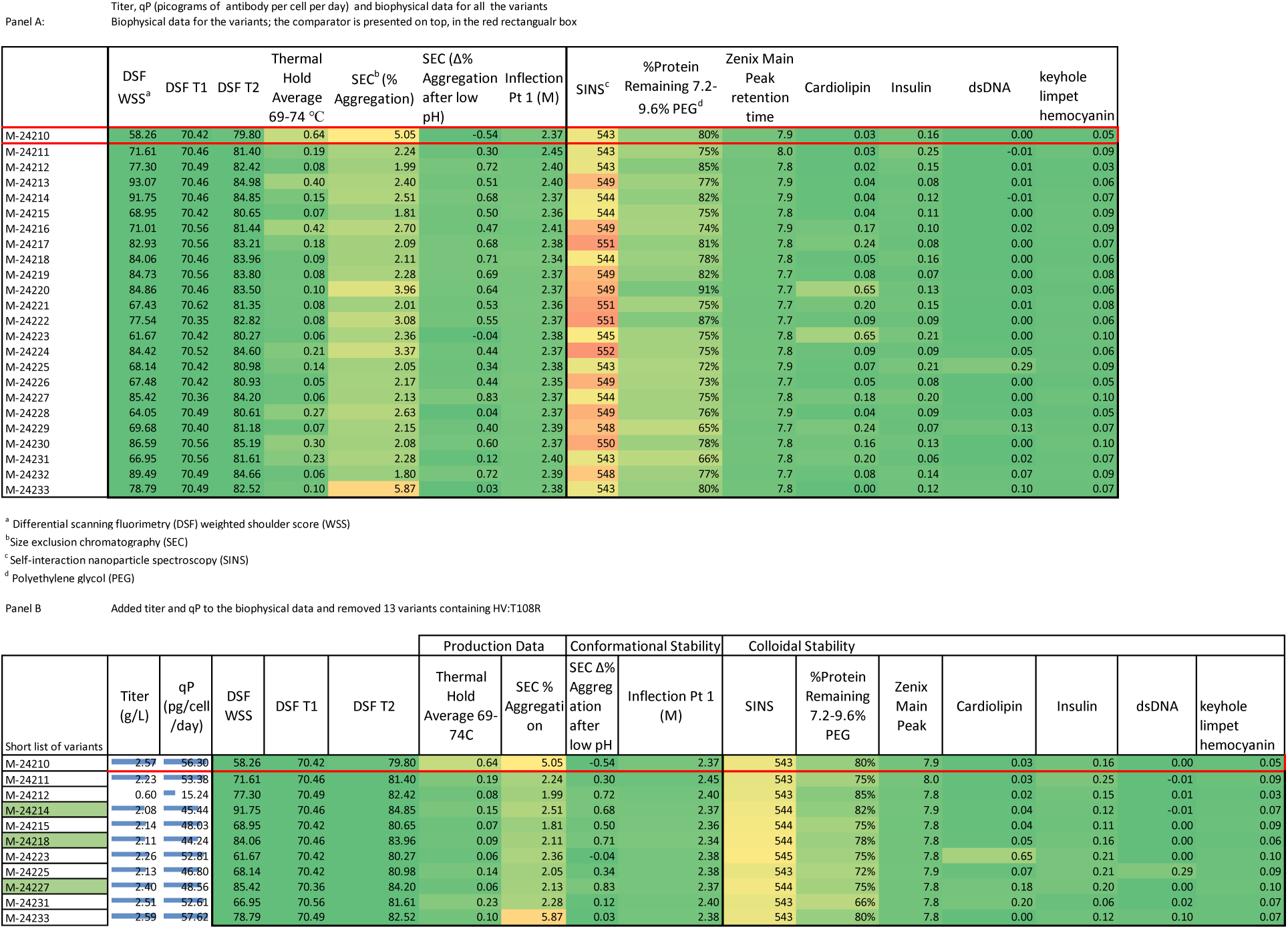
Titer, qP (picograms of antibody per cell per day) and biophysical data for all the variants.

### mAb Variants in Poliovirus Neutralization Assay

The 24 mAb variants were evaluated alongside the parental mAb 9H2 as a control in poliovirus neutralization assays. All variants demonstrated neutralizing activity against poliovirus type 1. For type 2, all variants were also neutralizing, with titration endpoints not reached, indicating high potency. All variants neutralized the Type 3 Sabin 3 strain with titers comparable to 9H2. For the neutralization titer against type 3 wild type capsid, S19 virus with type 3 capsid (S19 Saukett) had titers varied within approximately 3 log₂ across variants (Table 5) but remained comparable to the parental mAb. Overall, all variants retained neutralizing activity across all three serotypes, with titers similar to 9H2. Taken together, these data confirmed that the introduced sequence optimization was successful and preserved the broad and potent neutralizing capabilities of the variants relative to the parent mAb 9H2.

**Table 5:**
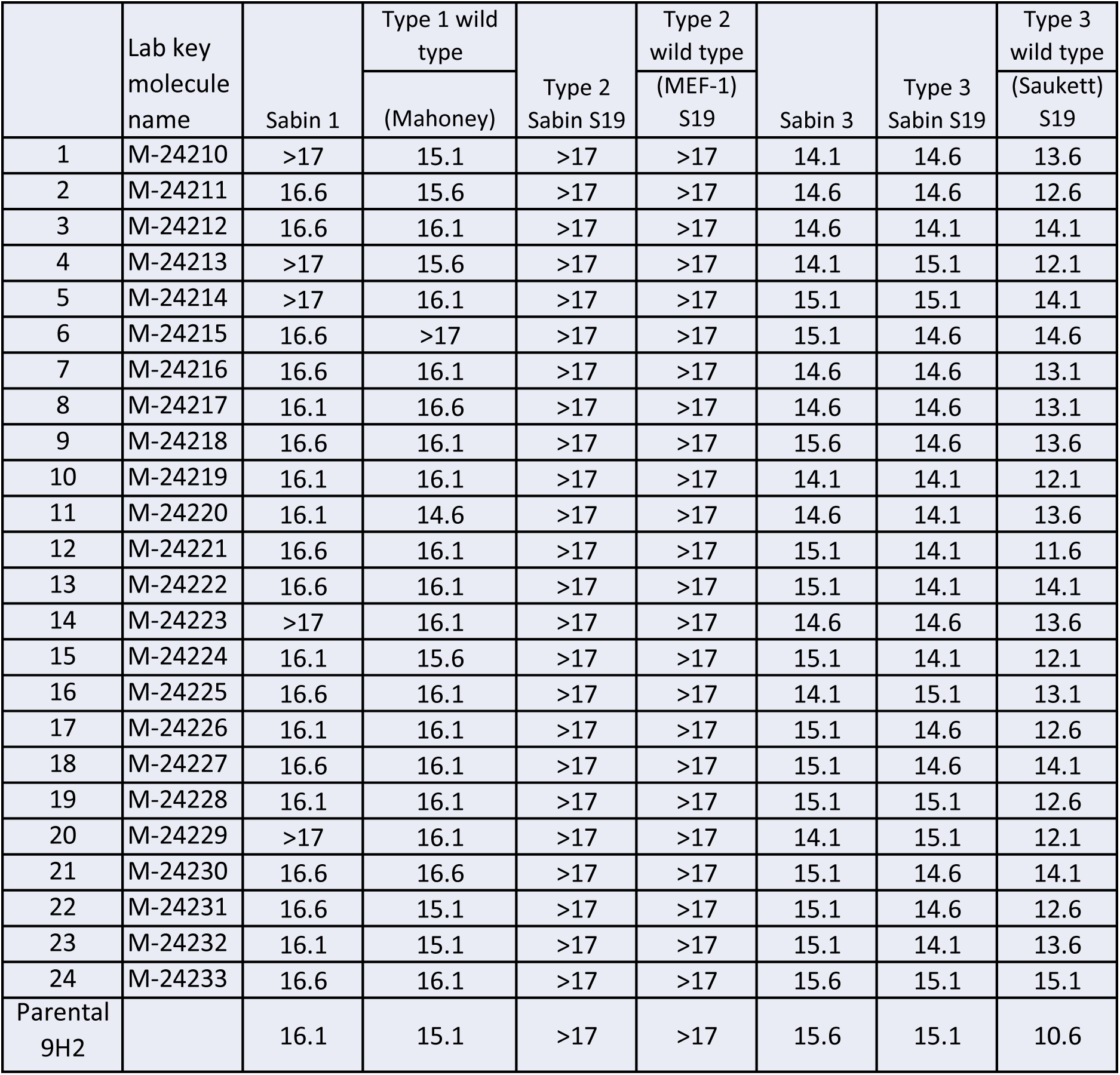
Log 2 of the neutralization activities for the 24 variants and parental 9H2.

|  | Lab key molecule name | Sabin 1 | Type 1 wild type | Type 2 Sabin S19 | Type 2 wild type | Sabin 3 | Type 3 Sabin S19 | Type 3 wild type |
| --- | --- | --- | --- | --- | --- | --- | --- | --- |
|  |  |  | (Mahoney) |  | (MEF-1) S19 |  |  | (Saukett) S19 |
| 1 | M-24210 | >17 | 15.1 | >17 | >17 | 14.1 | 14.6 | 13.6 |
| 2 | M-24211 | 16.6 | 15.6 | >17 | >17 | 14.6 | 14.6 | 12.6 |
| 3 | M-24212 | 16.6 | 16.1 | >17 | >17 | 14.6 | 14.1 | 14.1 |
| 4 | M-24213 | >17 | 15.6 | >17 | >17 | 14.1 | 15.1 | 12.1 |
| 5 | M-24214 | >17 | 16.1 | >17 | >17 | 15.1 | 15.1 | 14.1 |
| 6 | M-24215 | 16.6 | >17 | >17 | >17 | 15.1 | 14.6 | 14.6 |
| 7 | M-24216 | 16.6 | 16.1 | >17 | >17 | 14.6 | 14.6 | 13.1 |
| 8 | M-24217 | 16.1 | 16.6 | >17 | >17 | 14.6 | 14.6 | 13.1 |
| 9 | M-24218 | 16.6 | 16.1 | >17 | >17 | 15.6 | 14.6 | 13.6 |
| 10 | M-24219 | 16.1 | 16.1 | >17 | >17 | 14.1 | 14.1 | 12.1 |
| 11 | M-24220 | 16.1 | 14.6 | >17 | >17 | 14.6 | 14.1 | 13.6 |
| 12 | M-24221 | 16.6 | 16.1 | >17 | >17 | 15.1 | 14.1 | 11.6 |
| 13 | M-24222 | 16.6 | 16.1 | >17 | >17 | 15.1 | 14.1 | 14.1 |
| 14 | M-24223 | >17 | 16.1 | >17 | >17 | 14.6 | 14.6 | 13.6 |
| 15 | M-24224 | 16.1 | 15.6 | >17 | >17 | 15.1 | 14.1 | 12.1 |
| 16 | M-24225 | 16.6 | 16.1 | >17 | >17 | 14.1 | 15.1 | 13.1 |
| 17 | M-24226 | 16.1 | 16.1 | >17 | >17 | 15.1 | 14.6 | 12.6 |
| 18 | M-24227 | 16.6 | 16.1 | >17 | >17 | 15.1 | 14.6 | 14.1 |
| 19 | M-24228 | 16.1 | 16.1 | >17 | >17 | 15.1 | 15.1 | 12.6 |
| 20 | M-24229 | >17 | 16.1 | >17 | >17 | 14.1 | 15.1 | 12.1 |
| 21 | M-24230 | 16.6 | 16.6 | >17 | >17 | 15.1 | 14.6 | 14.1 |
| 22 | M-24231 | 16.6 | 15.1 | >17 | >17 | 15.1 | 14.6 | 12.6 |
| 23 | M-24232 | 16.1 | 15.1 | >17 | >17 | 15.1 | 14.1 | 13.6 |
| 24 | M-24233 | 16.6 | 16.1 | >17 | >17 | 15.6 | 15.1 | 15.1 |
| Parental 9H2 |  | 16.1 | 15.1 | >17 | >17 | 15.6 | 15.1 | 10.6 |

## Discussion

We have taken mAb 9H2 as a broad neutralizing mAb to all three poliovirus serotypes and analyzed it by MAM and Abacus™, identified its sequence liability, and designed sequence variants to extend its serum half-life and manufacturability. We further produced 24 sequence variants, compared them using eight biophysical methods, and downselected a top variant for future clinical production of a therapeutic anti-poliovirus mAb.

The design of 24 variants was successful; it did not alter the biological function of the antibody. All variants were tested to contain high neuralization titers to all three types of polioviruses, at the level comparable to parental 9H2. Remarkably, the remediation of the salt bridge disruption at heavy chain CDR3 position T108 was deleterious for the stability of the molecule, resulting in higher aggregation propensity. The remediation was suggested based on a strong deviation from germline distribution from the OAS at that position and common structural knowledge. This position, albeit peripheral, is still part of the CDR and the structure suggests that a mutation into an arginine might impact the nearby heavy chain CDR2, which is directly involved in antigen binding and, therefore, negates the benefits of restoring a salt bridge. Since determining a priori if the CDR3 salt bridge is required for binding is challenging, we opted to also include the original amino acid (threonine) for position HV_108 in the evaluation.

Although vaccination remains the central pillar of poliovirus prevention and global eradication, it has inherent limitations in the post-exposure context. IPV induces protective neutralizing antibodies but requires time for seroconversion and does not directly suppress ongoing viral replication in the gut. Once poliovirus has invaded the central nervous system, immunization cannot alter the course of paralytic disease. Furthermore, individuals with primary B-cell immunodeficiencies may mount inadequate humoral responses and can develop prolonged iVDPV infection with sustained viral shedding, representing a documented risk to eradication efforts [18]. These limitations emphasize the gap between preventive vaccination and direct antiviral intervention.

Accordingly, there is a strong scientific rationale for developing targeted antiviral therapeutics, particularly broadly neutralizing human monoclonal antibodies directed against conserved capsid epitopes. Structural and functional studies have identified mAbs capable of binding regions overlapping the poliovirus receptor (CD155) footprint and neutralizing multiple serotypes with high potency [7]. Such biologics have the potential to provide immediate passive immunity, rapidly reduce viral load and shedding, and serve as post-exposure prophylaxis or adjunctive therapy in high-risk populations. Integration of mAb-based interventions with vaccination strategies could therefore strengthen global polio control and support sustained eradication efforts.

In this context, our study demonstrates that a broadly neutralizing anti-poliovirus monoclonal antibody can be rationally engineered to improve developability while preserving potent cross-serotype activity, providing a clear path forward for monoclonal antibody–based therapeutics as a practical complement to vaccination in achieving and sustaining global polio eradication.

## Notes

## Acknowledgements

The authors thank the funding support from the Gates Foundation through grant to PATH (INV-010160). The authors also thank John Konz, Chris Gast, Steve Dong, Lauren Newhouse, Drew Conkin, and Sarah Calvillo for their contribution to the manuscript.

## Author contributions

S.-M.L.: Conceptualization, literature review, data curation, and writing the original draft.

C.S.: Methodology and formal analysis (MAM and biophysical data).

I.D.: Data curation (in silico analysis and variant design).

D.K.: Methodology, data curation, and formal analysis (neutralization assays).

K.M.: Conceptualization, supervision, and project administration.

All authors contributed to writing, review, editing, and approving the final manuscript.

## Disclaimer

The conclusions and opinions expressed in this work are those of the author(s) alone and shall not be attributed to the Gates Foundation. Under the grant conditions of the Foundation, a Creative Commons Attribution 4.0 License has already been assigned to the Author Accepted Manuscript version that might arise from this submission. Please note works submitted as a preprint have not undergone a peer review process.

